# Environmental correlates of genetic variation in the invasive and largely panmictic European starling in North America

**DOI:** 10.1101/643858

**Authors:** Natalie R. Hofmeister, Scott J. Werner, Irby J. Lovette

**Affiliations:** Department of Ecology and Evolutionary Biology, Cornell University, 215 Tower Road, Ithaca, NY 14853; Fuller Evolutionary Biology Program, Cornell Lab of Ornithology, Cornell University, 159 Sapsucker Woods Road, Ithaca, NY 14850; United States Department of Agriculture, Animal and Plant Health Inspection Service, Wildlife Services, National Wildlife Research Center, 4101 LaPorte Avenue, Fort Collins, CO 80521

**Keywords:** panmixia, gene flow, local adaptation, invasion genetics

## Abstract

Populations of invasive species that colonize and spread in novel environments may differentiate both through demographic processes and local selection. European starlings (*Sturnus vulgaris*) were introduced to New York in 1890 and subsequently spread throughout North America, becoming one of the most widespread and numerous bird species on the continent. Genome-wide comparisons across starling individuals and populations can identify demographic and/or selective factors that facilitated this rapid and successful expansion. We investigated patterns of genomic diversity and differentiation using reduced-representation genome sequencing (ddRADseq) of 17 winter-season starling populations. Consistent with this species’ high dispersal rate and rapid expansion history, we found low geographic differentiation and few F_ST_ outliers even at a continental scale. Despite starting from a founding population of approximately 180 individuals, North American starlings show only a moderate genetic bottleneck, and models suggest a dramatic increase in effective population size since introduction. In genotype-environment associations we found that ∼200 single-nucleotide polymorphisms are correlated with temperature and/or precipitation against a background of negligible genome- and range-wide divergence. Local adaptation in North American starlings may have evolved rapidly even in this wide-ranging and evolutionarily young population. This survey of genomic signatures of expansion in North American starlings is the most comprehensive to date and complements ongoing studies of world-wide local adaptation in these highly dispersive and invasive birds.

## INTRODUCTION

Studies of local adaptation have long bridged the interface between ecological and evolutionary questions by exploring how populations adapt to differing environmental conditions. Traditionally, high degrees of local adaptation were expected to be present only in fairly isolated populations—those free from the homogenizing effects of high gene flow—with a long history in those locations, providing the time thought to be necessary for local adaptation to evolve (Lenormand, 2002). We now know that local adaptation occurs frequently even in systems with high gene flow (Yeaman & Whitlock 2011; Tigano & Friesen, 2016) and often rapidly after colonization of a novel environment (Prentis *et al*., 2008). We continue to find evidence for rapid local adaptation in systems as divergent as cane toads (Rollins *et al*., 2015), sticklebacks (Lescak *et al*., 2015), honeybees (Avalos *et al*., 2017), steelhead trout (Willoughby *et al*., 2018), deer mice (Pfeifer *et al*., 2018), and many more. These studies show that many taxa can adapt rapidly to local conditions in response to the new selection regimes they encounter as they expand their range.

Invasive species that have recently expanded into new locations provide tractable opportunities to investigate local adaptation as it originates (Colautti & Lau, 2015). Invasions typically expand from the founding population(s) following a predictable spatial and temporal pattern (reviewed in Excoffier *et al*., 2009). After successful colonization of a new habitat, many invasive species show a demographic boom that may be facilitated by their ecological release in the new environment; ecological release is the concept that introduced species are often released from top-down limitations to their population growth, such as predators or pathogens in their native range (Riccardi *et al*., 2013). Theory predicts that this rapid population growth will plateau as the population approaches carrying capacity in a region, but successful invasive species may continue to expand their range and thus maintain a high rate of overall population growth.

When population density increases and demographic rates change, introduced species may rapidly evolve traits that enable them to spread (Szűcs *et al*., 2017): for example, invasive cane toads in Australia evolved a suite of morphological, physiological, and behavioral traits that facilitated their expansion in only 80 years (Rollins *et al*., 2015). Increased dispersal may evolve in concert with the demographic boom, such that certain traits are selected for in successful invasive species. Flexible dispersal strategies can result in gene flow that counteracts inbreeding depression and increases adaptive potential (Garant *et al*., 2007; Rius & Darling, 2014). If particular traits enable individuals to disperse more easily to their preferred habitat, gene flow may be directional and even adaptive (Edelaar & Bolnick, 2012; Jacob *et al*., 2017). Invasions thus allow us to observe interactions between demography and the early processes of selection (Dlugosch *et al*., 2015) as populations experience new environments. Importantly, these eco-evolutionary interactions depend in part on the genetic variation in the established population.

Recent work in invasion genetics aims to tease apart how selection and demography might resolve a paradox of invasion (Estoup *et al*., 2016, Schrieber & Lachmuth, 2017): many invasive species experience genetic bottlenecks as a result of an initial founder effect, but often thrive and spread despite this loss of standing genetic diversity. Theory predicts that introductions will typically result in an initial contraction in population size and/or genetic diversity (Dlugosch & Parker, 2008). However, bottlenecks of genetic variation clearly do not limit the success of many invasive species (Schmid-Hempel *et al*., 2007; Dlugosch & Parker, 2008; Facon *et al*., 2011). Invaders might adapt through soft sweeps that reduce genetic diversity while selecting for adaptive variants from standing or novel genetic variation, which is especially likely in the case of ecological adaptation (Messer & Petrov, 2013). Furthermore, some invasions may increase rather than reduce genetic diversity, as when multiple invasions from different source populations introduce previously isolated alleles and thereby facilitate admixture (Dlugosch & Parker, 2008). The new conditions can also select among standing variation, where the presence of certain genetic variants in the native range accelerates adaptation upon introduction (Tsutsui *et al*., 2000; Schlaepfer *et al*., 2009; Hufbauer *et al*., 2011). In sum, although genetic diversity in introduced populations is often viewed as a pre-requisite to adaptation, changes in genetic diversity alone do not explain invasiveness (Uller & Leimu, 2011). Regardless, chance may be just as important as selection in an invasive species’ establishment and spread (Gralka & Hallatschek, 2019).

The European starling (*Sturnus vulgaris*) stands out as an exceptionally successful avian colonist and invasive species. In North America, an estimated 200 million starlings currently range from northern Mexico to southern Alaska (Linz *et al*., 2007). Introduced to New York City in 1890, starlings nearly covered the continent within a few generations by expanding up to 91 km each year (Bitton & Graham, 2014). The 1890 introduction is widely accepted as the first successful establishment of starlings in North America, but several populations were introduced in Cincinnati, OH (1872), Quebec, Canada (1875), Allegheny, PA (1897), and Springfield, MA (1897), with the second-most successful having been introduced to Portland, OR in 1889 (Forbush, 1915; Kalmbach & Gabrielson, 1921). Records indicate that none of the earlier starling introductions survived more than a few years after colonization, but it is possible that some populations in the western U.S. persisted without record (Kessel, 1953).

During the starling expansion, ongoing migration and dispersal might have also influenced patterns of genetic variation. In North America, some—but not all—starling populations migrate (Dolbeer, 1982). Previous studies indicate that there is considerable variation in migratory distances within flyways (Burtt & Giltz, 1977). Models of molt origin indicate that starlings in the western U.S. may disperse or migrate shorter distances (Werner, Fischer, & Hobson, 2020). The models used feathers from the same individuals sampled for this genetic survey, and they initially hypothesized that starlings in the south may be less likely to migrate long distances. Instead, they found that longitude better explains the differences in molt origin among starling populations: starling feathers collected west of -90° longitude were more likely to have originated nearby. In other words, starlings that migrate in eastern North America likely experience greater gene flow among sampling locations (states, in our study), but overall starlings tend to move only regionally and not continent-wide.

Early genetic work based on a small set of allozyme markers indicated near-random mating at a continental scale in North American starlings, with large demes (subpopulations) and high dispersal rates (Cabe, 1998; Cabe, 1999). Here we use robust genomic markers to explore the genomic and demographic patterns of range expansion in North American starlings with three specific aims: (1) to characterize genome-wide levels of diversity and differentiation among starlings; (2) to examine how genetic variation changes across the contiguous United States; (3) to test for a genetic bottleneck; and (4) to test for signatures of selection associated with environmental gradients. We also interpret our results in the context of range-wide data on starling migration and dispersal (Werner, Fischer, & Hobson, 2020), as this movement certainly influences population structure. Recent developments in genotype-environment association methods can identify polygenic traits which are locally adapted to overlapping environmental gradients (Forester *et al*., 2018; Capblancq *et al*., 2018), whereas traditional outlier-based methods may not recover evidence of local adaptation in a species that likely has low overall levels of genetic diversity. Especially with these more sensitive methods, the same low genetic diversity resulting from a founder effect upon colonization could improve our ability to discriminate signatures of selection from low background divergence across the genome (Dlugosch *et al*., 2015). This work employs modern genomic and analytical tools to examine the evolutionary history of this remarkably successful avian invasion.

## METHODS

### Sample collection and processing

All starlings sampled were collected during the non-breeding season, when large flocks aggregate at high-quality food sources. These samples therefore do not represent discrete breeding populations but rather a mixture of birds from the surrounding region, potentially including migrants from more northerly areas. Starlings (*N* = 166) were collected in January-March 2016 and 2017 from 26 dairies and feedlots by the U.S. Department of Agriculture’s Wildlife Services personnel in Arizona, California, Colorado, Idaho, Illinois, Iowa, Kansas, Missouri, Nebraska, Nevada, New Hampshire, New Mexico, New York, North Carolina, Texas, Washington, and Wisconsin. USDA Wildlife Services personnel collected birds from 2-5 sites in each state, where each collection site was >5 km apart. USDA personnel then euthanized whole adult males by lethal gunshot, avicide, or live traps, stored at these birds at 0°C until tissue sampling, and recorded the latitude and longitude of each collection site using a handheld GPS. The collection and use of starlings for this and related studies were approved by the U.S. Department of Agriculture, National Wildlife Research Center’s Institutional Animal Care and Use Committee (QA-2572, S.J. Werner - Study Director; Werner, Fischer, & Hobson, 2020; Table S1).

Breast muscle tissue was sampled using biopsy punches (Integra Miltex, York, PA) and frozen in 95% ethanol. Samples were shipped on dry ice, and DNA was extracted using a Qiagen DNeasy kit following the manufacturer’s protocol (Qiagen, New York, NY). DNA concentration of each sample was quantified using a Qubit 2.0 fluorometer (Thermo Fisher Scientific, New York, NY). Following the protocol of Peterson *et al*. (2012), we generated a reduced-representation genomic dataset of double-digested, restriction-site associated DNA markers as described in Thrasher *et al*. (2017) using the restriction enzymes SbfI and MspI and adaptors P1 and P2. We sequenced 100bp, single-end reads on an Illumina HiSeq 2500. We trimmed and filtered for quality using the FASTX-Toolkit (http://hannonlab.cshl.edu/fastx_toolkit). We then used the process_radtags commands in _STACKS_ v 1.19 (Catchen *et al*., 2013) to demultiplex the remaining sequences. In subsequent filtering steps, we retained reads only if the following conditions were met: reads passed the Illumina chastity filter, contained an intact *Sbf*I RAD site, contained one of the unique barcodes, and did not contain Illumina indexing adaptors.

We assembled sequences to an *S. vulgaris* reference genome (Hofmeister, Rollins *et al*., *in prep*) using the ref-map option in _STACKS_. Individual reads were mapped to the reference genome using _BOWTIE_2 version 2.2.8 (Langmead & Salzberg, 2012) using the “very sensitive local” set of alignment pre-sets. The bioinformatics pipeline used for the reference-based assembly has the advantage of using fewer similarity thresholds to build loci. We required that a SNP be present in a minimum of 80% of the individuals (-r 0.8) with a minimum stack depth of 10 reads at a locus within an individual (-m 10) for it to be called. We removed two individuals, one with >50% missing data and one with >50% relatedness (measured using the unadjusted AJK statistic and calculated within *vcftools*), leaving 158 individuals remaining in the study. We also used *vcftools* –hwe option to remove any SNPs out of Hardy-Weinberg equilibrium (e.g., an exact test that compared expected and observed heterozygosity gave a P-value less than 0.001). About 6% of sequenced variants were out of HWE, and this filter left 14,134 variants remaining. For all genotype-environment analyses we used all SNPs in a given stack, but for STRUCTURE and other analyses sensitive to linkage disequilibrium we used only the first SNP in each stack; for unphased genomic data like the RAD loci analyzed here, this strategy of exporting one SNP per stack is often used as an indirect method of controlling for linkage.

#### (1) Patterns of genetic diversity and differentiation

We estimated per-locus measures of genetic diversity and genome-wide differentiation using the *populations* option in Stacks (Catchen *et al*., 2013). We used *vcftools* to calculate F_ST_ among population pairs and heterozygosity and nucleotide diversity (π) within populations (Danecek *et al*., 2011). We determined expected heterozygosity at the population level using the Hs() function in the R package *adegenet* (Jombart, 2008), and tested for differences in observed and expected heterozygosity of each locus using a paired t-test in base R (R Core Team, 2019). We investigated genetic structure within North American starlings using an analysis of molecular variance (AMOVA) in the R package *poppr* (Kamvar *et al*., 2014). We tested whether most genetic variation was observed among individuals or among sampling sites (“populations”). To determine significance, we compared observed variation at each hierarchical level to the randomly permuted distance matrices for that particular level using 1000 simulations in the function *randtest()* in the R package *adegenet* (Jombart, 2008), hypothesizing that the observed variance is greater than expected within individuals and less between individuals and between populations.

We tested for isolation by distance (IBD) using a simple Mantel test in *adegenet* (Jombart, 2008): for these data, the assumption of stationarity likely holds, given that North American starlings appear to be in mutation-migration-drift equilibrium (Guillot & Rousset, 2013). Geographic distances among sampling locations follow a bimodal distribution where locations are either very near or far from each other, and thus we caution that the Mantel test may not be an appropriate test of isolation by distance (Supplementary Information). Nevertheless, we report the results of a Mantel test with a Spearman correlation and 10000 permutations. Finally, we test for isolation by environment (IBE) using the R package *vegan* to compare environmental, geographic, and genetic distances (Oksanen *et al*., 2019). Controlling for geographic distance, we ran partial Mantel correlograms using Euclidean environmental distance matrices built from all bioclimatic variables retained in the *Selection analyses* described in Section 4 below.

#### (2) Population structure

We first used non-parametric approaches to determine whether starlings clustered by sampling location, using principal components analysis in *SNPRelate* (Zheng *et al*., 2012) and discriminant analysis of principal components (DAPC) in *adegenet* (Jombart, 2008). We then tested for population structure using STRUCTURE (Pritchard *et al*., 2000) by simulating 10 runs of each *K*. Although we sampled starlings from 17 locations, we hypothesized that North American starlings would cluster into at most eight populations (*K*=1-8) based on the non-parametric tests described above. To select the best-supported K, we used the Evanno method implemented in STRUCTURE HARVESTER v0.6.94 (Earl & vonHoldt, 2011). We averaged results across the 10 runs using the greedy algorithm in the program CLUMPP v1.1.2 (Jakobsson & Rosenberg, 2007), and visualized results using DISTRUCT v1.1 (Rosenberg, 2003). Given that evidence of population structure depends on the filtering thresholds selected, we ran this model-based approach using a very strict minimum minor allele frequencies (MAF=0.3) and a more relaxed minimum frequency (MAF=0.1) (Linck & Battey, 2017). STRUCTURE results did not differ substantially among MAF thresholds, and we used a filtered dataset with a minimum minor allele frequency of 0.1 in the demographic analyses below.

#### (3) Demographic modeling

We used a traditional model-based approach (fastsimcoal2) to explicitly test for a bottleneck in North American starlings (Excoffier *et al*., 2013). We first estimated the folded SFS using a Python script from Simon Martin (available at https://github.com/simonhmartin/genomics_general). We modified the simple 1PopBot20Mb.tpl and .est files provided with the software as follows: the size of the bottleneck (NBOT: 10-1000), start of bottleneck (TBOT: 10-300), ancestral size (ANC: 1000-100000), current size of the population (NCUR: 100-100000), and end of bottleneck (TENDBOT: TBOT + 500). We performed 100 runs of this model and assessed model fit using delta-likelihood for each run of the model; since all models use the same numbers of parameters, the Akaike information criteria do not change between models. We also ran a demographic model that takes linkage into account, the Stairway plot method, but given that this method does not explicitly test for a genetic bottleneck, we describe the method and results in the Supplementary Information.

#### (4) Selection analyses

We first identified environmental variation at each sampling location using the R package raster (Hijmans & van Etten, 2012), extracting all 19 bioclimatic variables for each set of sampling coordinates from WorldClim 2.0 at a resolution of 5 min on June 16, 2018 (Fick & Hijmans, 2017). We recognize that these environmental conditions represent those experienced during the non-breeding season, and therefore they do not represent the full range of conditions experienced by the starlings sampled; see the Supplementary Information for a discussion of our assumptions about this environmental sampling. We used redundancy analysis (RDA) to examine how loci may covary across multiple environmental gradients (Forester *et al*., 2018). RDA is especially powerful when testing for weak selection, and detects true positives in large data sets more reliably than other multivariate methods like Random Forest (Forester *et al*., 2018). Because RDA requires no missing data, we first imputed genotypes where missing sites were assigned the genotype of highest probability across all individuals—a conservative but quick imputation method. Before imputation and after filtering, about 31% of genotypes were missing from the dataset of 15,038 SNPs (not filtered for MAF), and we required that all imputed SNPs were present in at least 80% of individuals. We then used the R package *vegan* to run the RDA model (Oksanen *et al*., 2019); for a full description of this method, see Forester *et al*., 2018. Briefly, RDA uses constrained ordination to model a set of predictor variables, and unconstrained ordination axes to model the response (genetic variation). RDA infers selection on a particular locus when it loads heavily onto one or more unconstrained predictor axes, and we retained five variables with relatively low variance inflation factors: BIO1 or Annual Mean Temperature (VIF=3.54), BIO7 or Temperature Annual Range (4.55), BIO12 or Annual Precipitation (8.69), BIO16 or Precipitation of Wettest Quarter (7.91), and elevation (2.19). We tested for significance using the *anova*.*cca* function within the *vegan* package, and also permuted predictor values across individuals to further check significance of the model. We identify candidate loci as those that are 3 or more standard deviations outside the mean loading. The R scripts for all RDA analyses and figures were written by Brenna Forester (available at https://popgen.nescent.org/2018-03-27_RDA_GEA.html).

## RESULTS

### (1) Patterns of genetic diversity and differentiation

We identified 15,038 SNPs at a mean of 27X sequencing depth across 17 sampling locations. Using a dataset filtered by a minor allele frequency of 0.1, we find that genome-wide F_ST_ is extremely low (0.0085), and measures of genetic diversity do not vary substantially among sampling locations (Table 1). Across all of these wintering starling populations, the highest pairwise F_ST_ was 0.0106 between birds from the adjacent states of Arizona and New Mexico. Using a haplotype-based statistic of differentiation, ϕ_ST_ among populations shows an absence of genetic structure (ϕ_ST_ = 0.0002). Hierarchical AMOVAs reveal that 94% of the observed genetic variance is explained by variation within individuals, and the remaining variance reflects differences among individuals in the same population, with negligible variation explained at the between-population level (Figure 1C-D). Across the genome, F_ST_ and nucleotide diversity are exceptionally low (Table 1,2). Genome-wide heterozygosity is moderate at 0.339, and across loci observed heterozygosity differs significantly from expected (t = 66.6, df = 3569, P<0.001, Table 1).

**Figure 1.**
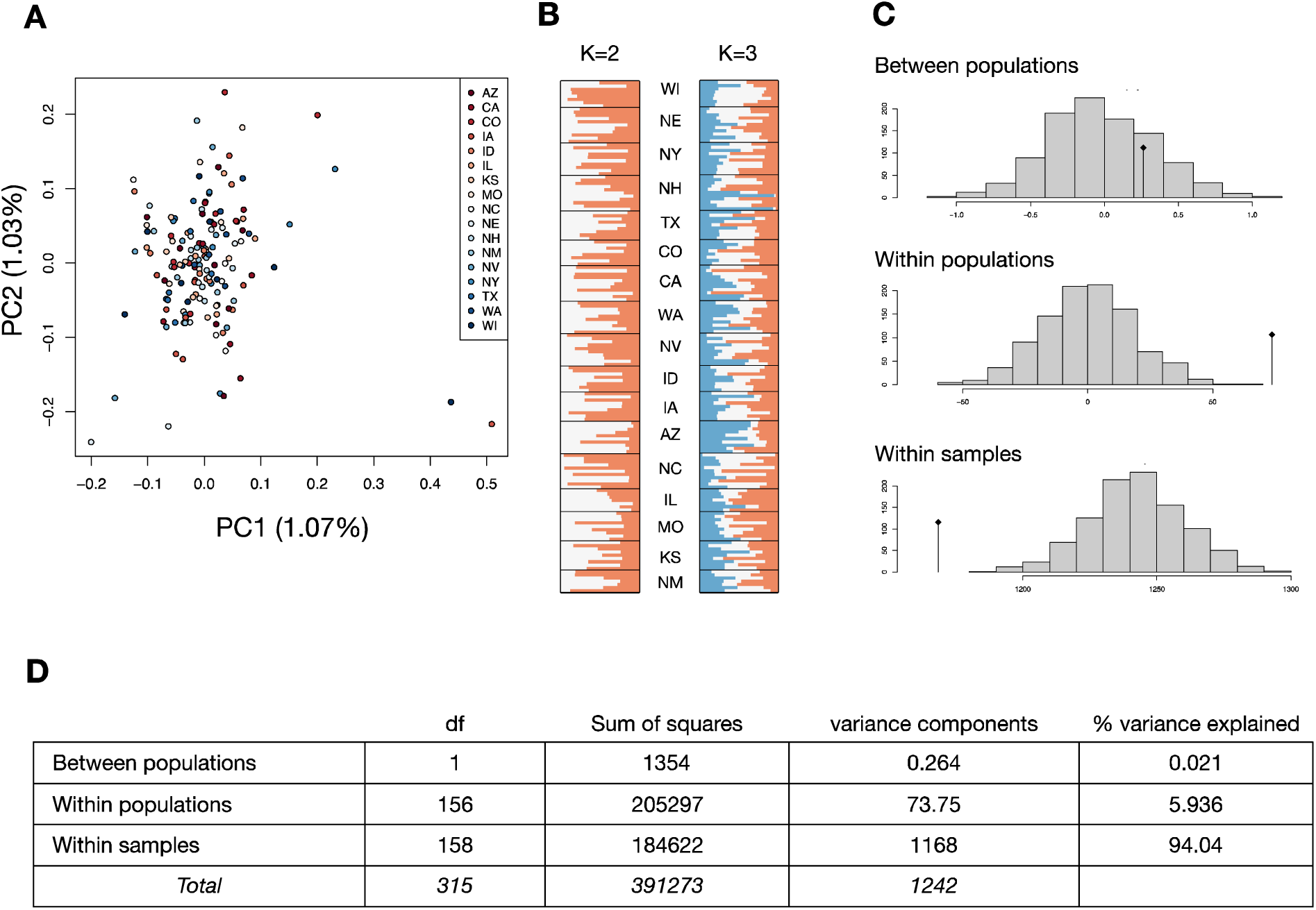
Population structure. A) Principal components analysis on 6,287 SNPs, explaining 1.03-1.08% of genetic variation observed. B) STRUCTURE analyses with K=2 and K=3 (best supported). C) Significance testing of hierarchical AMOVAs: the histogram shows expected variance components based on 999 simulations, and the black diamond is the observed variance component. D) AMOVA results.

**Table 1.** Population genetic summary statistics. For each sampling location, the table shows the number of private alleles, observed and expected heterozygosity, nucleotide diversity, and the inbreeding coefficient F_IS_.

| Pop. | No. | Private | H<sub>obs</sub> | H<sub>exp</sub> | $\pi$ | F<sub>IS</sub> |
| --- | --- | --- | --- | --- | --- | --- |
| NY | 10 | 0 | 0.34 | 0.25 | 0.35 | 0.037 |
| NH | 11 | 1 | 0.33 | 0.26 | 0.35 | 0.065 |
| NC | 11 | 1 | 0.33 | 0.25 | 0.35 | 0.060 |
| IL | 7 | 1 | 0.33 | 0.26 | 0.35 | 0.056 |
| IA | 9 | 0 | 0.33 | 0.24 | 0.36 | 0.078 |
| WI | 8 | 0 | 0.33 | 0.22 | 0.35 | 0.050 |
| MO | 9 | 0 | 0.33 | 0.23 | 0.35 | 0.053 |
| NE | 11 | 0 | 0.34 | 0.22 | 0.35 | 0.042 |
| CO | 8 | 1 | 0.33 | 0.24 | 0.35 | 0.066 |
| KS | 9 | 2 | 0.35 | 0.23 | 0.35 | 0.012 |
| TX | 9 | 1 | 0.33 | 0.24 | 0.35 | 0.052 |
| NM | 7 | 2 | 0.33 | 0.29 | 0.36 | 0.068 |
| AZ | 10 | 1 | 0.32 | 0.24 | 0.35 | 0.069 |
| NV | 10 | 0 | 0.33 | 0.31 | 0.36 | 0.068 |
| CA | 11 | 0 | 0.33 | 0.25 | 0.35 | 0.061 |
| ID | 8 | 0 | 0.33 | 0.22 | 0.35 | 0.047 |
| WA | 10 | 2 | 0.32 | 0.30 | 0.35 | 0.082 |

### (2) Population structure

F_ST_ among populations is relatively low overall (Table 2): starlings sampled in Arizona show the highest differentiation from other sampling locations (F_ST_ = 0.008-0.011), and only a few other pairwise comparisons (NM-IL, NM-CO, and CO-WI) show an F_ST_ of 0.009 or higher. However, we do not recover clear population structure. The first two principal components each explain about 1% of variation among individuals (Figure 1A), and although STRUCTURE identified three populations at the best-supported value of K, these predicted populations do not show obvious differences in ancestry proportions (Figure 1B, Figure S1). Controlling for shared ancestry does not resolve population structure, and instead provides support for uniform gene flow among individuals (Figure S2). K-means clustering within DAPC also does not identify biologically relevant clusters.

**Table 2.**
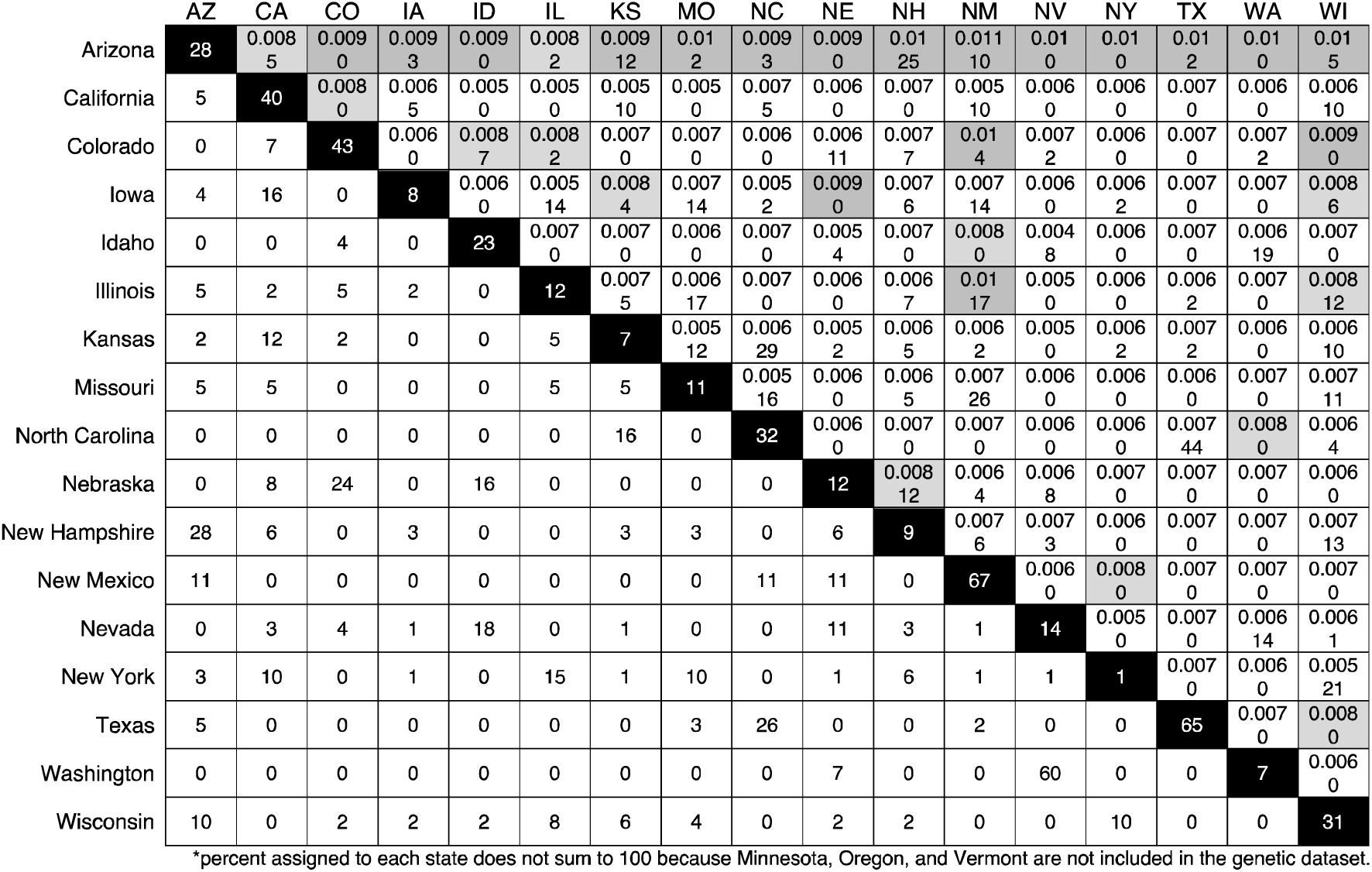
Differentiation and movement among sampling locations. This table shows the percentage of birds assigned to each sampling location from every origin. Note that these values are directional: although 12 starlings that were collected in AZ originated in KS, only 2 starlings collected in KS originated in AZ. The black diagonal indicates the percentage of birds assigned to that collection state according to a discriminant function analysis of molt-origin presented in Werner *et al*. For example, 28% of birds collected in Arizona originated in that location. F_ST_ among locations is presented above the diagonal, and darker gray colors indicate higher F_ST_.

Starling populations follow a clear pattern of isolation-by-distance (Mantel *r* = 0.139, P < 0.0001). Spatially explicit models of isolation-by-distance suggest a fairly uniform rate of migration range-wide, where local increases in migration rate are likely a model artifact (Supplementary Information). However, models of isolation by environment (IBE) show that the relationship between environmental and genetic distances is stronger than the simple geographic-genetic distance model. After controlling for geographic distance, we find that all bioclimatic variables tested show non-zero correlations between environmental and genetic distances (Supplementary Information). There is a strong positive relationship between genetic distance and both precipitation in the wettest quarter (BIO16, Mantel *r* = 0.282, P = 0.001) and elevation (Mantel *r* = 0.146, P = 0.001).

### (3) Demographic modeling

A traditional SFS-based model recovers a bottleneck in population size, as expected: the estimated effective population size during the bottleneck is ∼16,000 individuals (1% of the ancestral population size). However, our demographic model suggests that the current population size of North American starlings is nearly identical to its pre-bottleneck size (Supplementary Information). In fact, every run of the model finds that starlings’ current effective population size (1.6 million individuals) is considerably higher than the estimated N_e_ of the founding population. This model suggests that starlings experienced rapid population growth after the initial founder effect, which may contribute to the overall lack of evidence for inbreeding. We do detect very low levels of inbreeding within some populations (Table 1, highest F_IS_ = 0.082 in Washington). Taken together, these models do not suggest a classical founder effect, whereby effective population size remains very low for many generations post-bottleneck.

### (4) Selection analyses

Starlings encounter a range of precipitation, temperature, and elevation across their range and redundancy analyses revealed the strongest signatures of local adaptation, showing that 178 variants are correlated with environmental differences among populations (F = 1.022, P = 0.002, Figure 2). Populations living in warmer climates tend to cluster more closely in the left quadrant and high elevation populations cluster in the middle right quadrant. However, populations do not cluster based on geographic distance: for example, starlings from TX and WA cluster closely due to shared genetic variants, even though the two populations differ substantially in precipitation and temperature and are geographically separated. After controlling for population structure, candidates for selection are equally distributed among elevation, temperature- and precipitation-associated predictors. Importantly, when environmental predictors are randomly shuffled, the RDA model is not significant. Mean annual temperature (BIO1) opposes selective pressure related to the range of temperatures experienced each year (BIO7), annual precipitation (BIO12), precipitation in the wettest quarter (BIO16) and elevation (Figure 2). Genes under selection tend to have lower allele frequencies, and function in several biological processes (Table S2).

**Figure 2.**
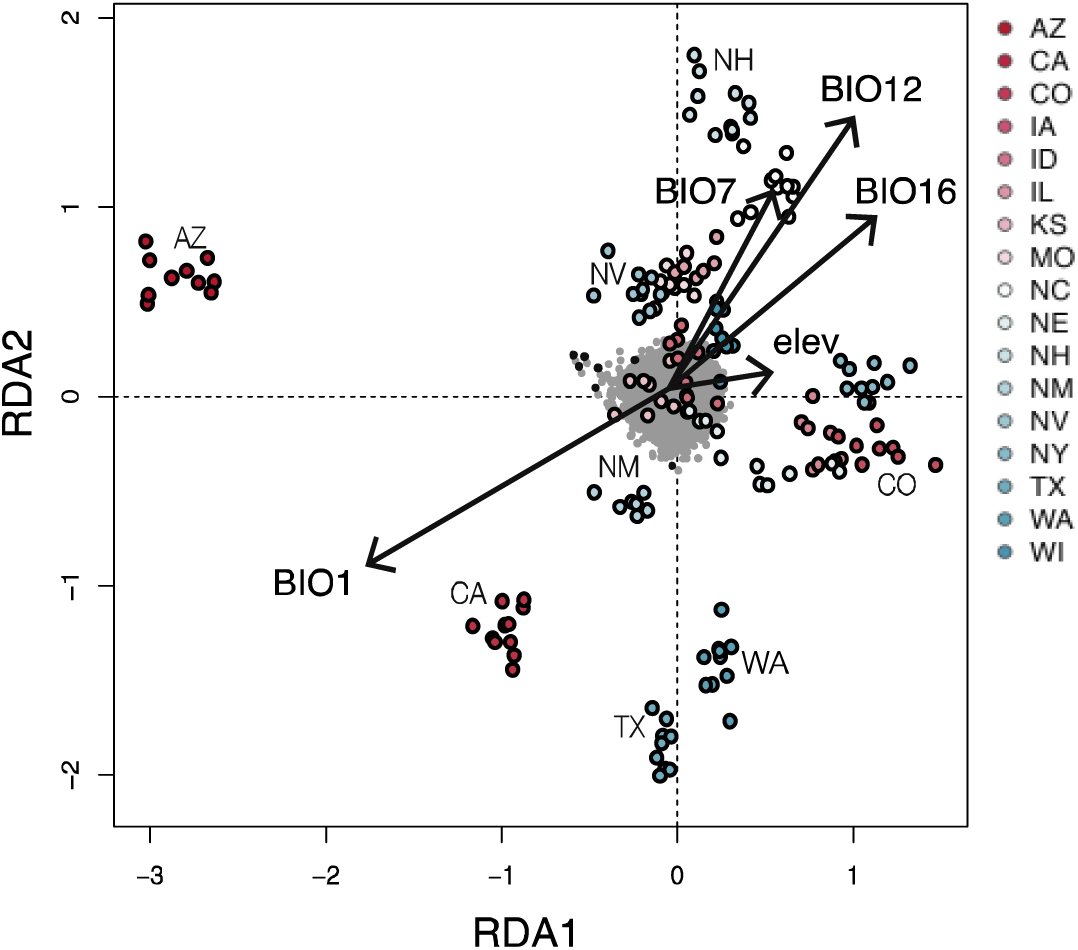
Evidence for incipient local adaptation. Redundancy analyses indicate that 191 SNPs (small gray points) are associated with bioclimatic predictors (vectors). BIO1: mean annual temperature; BIO7: temperature annual range; BIO12: annual precipitation; BIO16: precipitation of wettest quarter.

## DISCUSSION

Our whole-genome data reveal that invasive starling populations show moderate genetic diversity across North America, rapid population growth after a genetic bottleneck, negligible geographically structured genetic differentiation, significant but quite weak isolation-by-distance, and intriguing associations of genetic variation with environmental parameters that might result from adaptive processes. Genetic diversity is typically predicted to be low in invasive populations (Dlugosch & Parker, 2008), and although RADseq tends to estimate lower genetic diversity than the true diversity, this bias is minimal for taxa known to be genetically depauperate (Cabe, 1998; Cariou *et al*., 2016). Within invasion genetics, one area of recent focus involves the spatial dynamics of invasion, and we can explore how genetic variation varies across the post-invasion range of starlings in North America. Although there is a significant (but low magnitude) signal of isolation-by-distance, hierarchical AMOVAs find that variation within and among individuals explains observed differentiation better than variation among populations. There is no evidence of population structure, and while models indicate some subtle spatial patterns of genetic variation, these model-based inferences likely reflect sampling artifacts (Supplementary Information). Finally, after the initial founder effect, the effective population size has grown by a ratio of 1:100. These patterns are consistent with the expectation that extensive gene flow—as shown by extremely low F_ST_ among populations (Table 2)—maintains high connectivity across North American starling populations. When interpreted within the context of a complementary exploration of movements inferred from feather isotope assays, this genomic survey confirms that dispersal and migration continue to influence the genetic variation generated in just over a century of range expansion.

Long-distance dispersal may be common in introduced starlings on multiple continents, where rates of dispersal are determined by demographic changes and environmental quality. In South Africa, invasive starlings disperse when their natal environment becomes crowded or unsuitable, as at the leading edge of their range expansion (Hui *et al*., 2012). In North America, the effective population size (N_e_) of the present-day invasive population has expanded dramatically, with models indicating that current N_e_ is even larger than the N_e_ of the founding population. This explosive population growth from hundreds to millions of individuals may have encouraged long-distance dispersal away from the dense populations in eastern North America. In general, multiple lines of evidence have shown that the dispersal rates and distances of juvenile starlings in North America are remarkably high (Dolbeer, 1982; Cabe, 1999; Werner, Fischer, & Hobson, 2020). Considered in concert with similar findings in the invasive South African and Australian starling populations (Hui *et al*., 2012; Phair *et al*., 2017), it is clear that dispersal similarly plays an ongoing and major role in the connectivity of North American starling populations.

Although it is generally important, dispersal has not been uniform across time or space in North American starlings. One open question in invasion genetics is whether expanding populations maximize spatiotemporal fitness via spatial sorting and incipient adaptation (Williams *et al*. 2019). Empirical studies of some invasions suggest that spatial sorting is common, including in common mynas (Berthouly-Salazar *et al*., 2012), pumpkinseed fish (Ashenden *et al*., 2017), and cane toads (Brown *et al*., 2014). More empirical work is needed to verify the conditions in which spatial sorting can lead to lasting shifts in fitness (Lee, 2011; Phillips & Perkins, 2019; Williams *et al*., 2019). However, we do know that adaptive dispersal strategies can facilitate range expansion in Western bluebirds (Duckworth, 2008) and invasive beetles (Lombaert *et al*., 2014; Ochocki & Miller, 2017) among other species. For North American starlings, we can combine historical records, isotopic and genetic evidence to make inferences about the possibility of spatial sorting.

Feather isotope data from the same starlings sampled in this genetic survey suggests that movements are not uniform continent-wide, and that both juvenile dispersal and regional differences in migration could influence genetic variation (Werner, Fischer, & Hobson, 2020). If starling movement was the primary driver of differentiation, then we would expect F_ST_ among populations to be directly related to the number of birds collected and assigned to that population. Starlings in the western U.S. appear to have differentiated subtly from their eastern counterparts based on the higher F_ST_ between Arizona—and to some degree, New Mexico and Colorado—and all other sampling locations (Table 2). Birds collected in these southwestern states are also assigned to those states by discriminant function analysis (Werner, Fischer, & Hobson, 2020); for example, 67% of starlings collected in New Mexico were also assigned to that state (Table 2). We suggest that birds assigned to the same state where they were collected may reside in that state year-round, but we note that collecting feathers once in the lifespan of the bird does not allow us to determine the bird’s lifelong migration and dispersal. The highest rates of pairwise differentiation occur between states where one of those “populations” shows a relatively high collection-state assignment, which suggests that starling movement among sampling locations does impact the genetic variation. This comparison is further supported by the low but significant isolation-by-distance across the North American range; however, environmental pressures explain genetic variation better than a simple model that accounts for geographic distance. In addition, assignment to a state other than the collection state seems to have only a minor effect on F_ST_, even though we might expect that movement among geographically distant sampling locations might lead to gene flow among those locations. In Arizona—where F_ST_ is highest but still relatively weak—Werner *et al*.’s model suggests that 25% of birds originated in New Hampshire, which is approximately the percentage of birds assigned to Arizona (28%, Table 2). Unsurprisingly, geography, environment, and demography all appear to influence genetic variation within North American starling, and explicit tests for selection can help to clarify the relative weight of each factor.

This species’ general genome-wide panmixia across the continent allows for tests of selection on loci that may be involved in local adaptation. Because genetic diversity and differentiation are exceptionally low, it is relatively easy to identify sites that have differentiated against this background of low genome-wide or population-specific divergence (Dlugosch *et al*., 2015). We find that almost 200 of the 15,038 RAD loci appear to be under selection using a redundancy analysis. Only 13 of these SNPs overlap with the SNPs identified by a latent-factor mixed model, and there is no overlap between the RDA candidates and the differentiation-based scans (BayeScEnv and Bayescan). It is unsurprising that each test identifies different candidates for selection, because the assumptions underlying each are very different (for more details on our approach to selection testing, see Supplementary Information). Rather than making inferences based on the genes identified by these scans, we instead propose that genotype-environment associations show that changes in precipitation and temperature can explain genetic variation in North American starlings. However, we do suggest that rapid evolution in genes underlying key physiological processes may have supported the starling’s spread across North America (Supplemental Information). Specifically, we hypothesize that aridity and cold temperatures that are not experienced in the starling’s native range exert enough selective pressure on North American starlings to result in incipient local adaptation. While this finding suggests that local adaptation may explain genetic variation within the North American starling invasion, there are several relevant caveats to this approach.

Because there is evidence of a genetic bottleneck in the North American starling invasion, it is possible that artifacts resulting from genetic drift could mimic these patterns of local adaptation. We cannot rule out allele surfing during range expansion as an explanation based on the methods currently available for testing genotype-environment associations in a young and expanding system like this one. However, given how short the timescale of divergence is in North American starlings, such drift effects are more likely if genetic variation is (1) ancestral and (2) structured among populations. Gene flow among starling populations is remarkably high, but we do not know if variants putatively under selection were present in the ancestral population. These alleles under putative selection do not approach fixation, as no putative outlier has an allele frequency greater than 0.28. We note that concordance among environmental and genetic distances (e.g., partial Mantel tests) indicate that spatial autocorrelation complicates our selection inferences. Under these conditions, any evidence for selection is likely to be weak, and as always these selection scans can generate false positives. However, RDA has the highest rate of true positives and lowest of false positives, and although this method has not been tested in such recent expansions, RDA is well-suited to systems where F_ST_ is very low (Meirmans, 2015, Forester *et al*., 2018, Supplementary Information). Finally, we do not explicitly control for linkage, and some allele frequency shifts could be explained by recombination or by genetic hitchhiking.

Our study focuses on birds collected during the winter, which may limit our inferences about population structure and selection to these wintering populations. As discussed above, isotopic evidence suggests that starlings in the western U.S. tend to move only regionally whereas birds sampled in the eastern U.S. undertake longer movements (Table 2). This in turn suggests that starlings overwintering in the western U.S. are more likely to breed nearby, and thus the environmental conditions may not change as dramatically among wintering and breeding ranges. In addition, the environmental conditions that we expect to drive selection—precipitation and temperature—vary most substantially in the southwestern region: for example, the sampling location in Arizona is consistently warmer (BIO1) and drier (BIO12 and BIO16) than other locations (Figure 2, Supplementary Information). Western populations experience these environmental conditions year-round, which could allow selection to drive advantageous alleles toward fixation. Elsewhere in the U.S., starlings move more freely among states: individuals within each sampling location may come from different breeding populations, and additional sampling could reveal stronger population structure among true breeding populations. However, if our sampling overlooked some true populations, we would expect some signal of population structure. Individual-based tests of population structure—e.g., those that do not define possible populations *a priori*—do not recover any signals of population structure. This sampling strategy uses the more vagile eastern populations as a comparison to more-resident western populations that may also be under stronger selection. This framing may suggest that population structure in the western U.S.—of which we find no evidence—could explain allele frequency shifts that we infer to be selection, but when we compare the relative importance of geographic and environmental distances in partial Mantel tests, we find that environmental conditions better explain genetic variation. This evidence supports our interpretation of selection as a major driver of genetic variation in North American starlings.

A similar project on starlings in the Australian invasion—which colonized that continent nearly concurrently with the North American invasion—found that geographic but not environmental distance explains genetic patterns there (Cardilini *et al*., 2020). Starlings in the Australian range show substantial population structuring and significant patterns of isolation-by-distance. Earlier work had shown that gene flow among Australian starling populations is low (Rollins *et al*., 2009), and phylogeographic patterns of mitochondrial sequence variation confirm that starlings on the edge of the expansion front in Western Australia have differentiated from those still living in the introduction site (Rollins *et al*., 2011). In fact, starlings at the expansion front may have rapidly adapted during the Australian invasion (Rollins *et al*., 2016): the proportion of adult starlings in Western Australia carrying a novel mitochondrial haplotype has increased rapidly only at this range edge. A genotyping-by-sequencing survey employing a much greater number of SNP markers indicates three population subdivisions in Australia, where geographic distance explains genetic differentiation in starlings better than does environmental variation (Cardilini *et al*., 2020). Global F_ST_ across all Australian populations is an order of magnitude higher than the equivalent F_ST_ index across North America, despite similar areas sampled. In Australia, the strong evidence for isolation-by-distance and founder effects complicate attempts to disentangle selection from drift, yet despite their differences in invasion dynamics, genotype-environment associations reveal signatures of selection in both invasions. On both continents, starling genetic variation can be explained by extremes in temperature and precipitation, and preliminary results of whole-genome resequencing of native and introduced populations confirm that variability in temperature and precipitation may shape observed genetic variation in starlings world-wide (Hofmeister et al., *in prep*).

Our results contribute to the growing evidence of rapid adaptation in some expanding populations, even in extremely young systems. Some studies of rapidly expanding invasions find little evidence that adaptation may facilitate this expansion, as in corals (Leydet et al. 2018). However, other work suggests a role for selection in supporting rapid range expansion, such as in experimental studies of flour beetles (Szucs *et al*., 2017) and empirical work in guppies (Baltazar-Soares *et al*., 2019). Invasion biologists have long highlighted propagule pressure as a driver of invasion success, but the genetic composition may be just as important as the size of the establishing population (Briski *et al*., 2018). For example, genetic bottlenecks in monk parakeets, another avian invader now distributed world-wide, do not seem to inhibit invasion success (Edelaar *et al*., 2015). Pre-adaptation in the native range or selection during transport may facilitate the spread of invasive species, and human commensalism may support establishment and spread, as shown in house sparrows (Ravinet *et al*., 2018) and common mynas (Cohen *et al*., 2019), and reviewed across alien bird species (Cardador & Blackburn, 2019). Empirical studies of invaders like the ones described here also show how, in addition to genetic variation, epigenetic shifts and/or plastic changes in gene expression may support the establishment and expansion of invasive species (Marin *et al*., 2019). In the well-studied house sparrow— a system quite similar to starlings—epigenetic shifts may have supported invasions in Africa (Liebl *et al*., 2013) but not necessarily in Australia (Sheldon *et al*., 2018). Taken together, recent work suggests that we should consider a much wider range of demographic and ecological processes that lead to adaptive evolution in invading populations.

Invasive populations allow us to explore the genetic consequences of colonization and establishment in novel environments. On a background of low genetic differentiation and diversity, we find evidence of incipient genotype-environment associations in North American starlings. Here we explore how genetic variation changes across the landscape, but we cannot fully understand gene flow without studies of dispersal and migration of the individuals that carry genes. Our results complement other recent studies that reveal associations between climate variables and particular loci in North American vertebrates (Schweizer *et al*., 2015; Bay *et al*., 2018). Finally, we suggest that our study adds to those suggesting that rapid local adaptation can evolve even in dispersive and young populations.

## Supporting information

SI: BayeScEnv Results

SI: Demography

## AUTHOR CONTRIBUTIONS

NRH, IJL designed research, SJW contributed samples, NRH analyzed data, NRH, IJL wrote paper.

## ACKNOWLEDGEMENTS

We are grateful to the U.S. Department of Agriculture’s Wildlife Services personnel in Arizona, California, Colorado, Idaho, Illinois, Iowa, Kansas, Missouri, Nebraska, Nevada, New Hampshire, New Mexico, New York, North Carolina, Texas, Washington, and Wisconsin, who collected samples used here. Jennifer Walsh-Emond, Leonardo Campagna, Daniel Hooper, Jacob Berv, and Stepfanie Aguillon provided valuable advice on bioinformatic methods.

## DATA ACCESSIBILITY STATEMENT

All scripts are archived on GitHub: https://github.com/nathofme/radseq-NAm All data files that accompany the above scripts are archived on Dryad (doi: 10.5061/dryad.j424f07), and raw sequencing data archived in NCBI SRA (Bioproject PRJNA545151).

