## Supplementary Text for "Environmental correlates of genetic variation in the invasive and largely panmictic European starling in North America"

### Tests of population structure

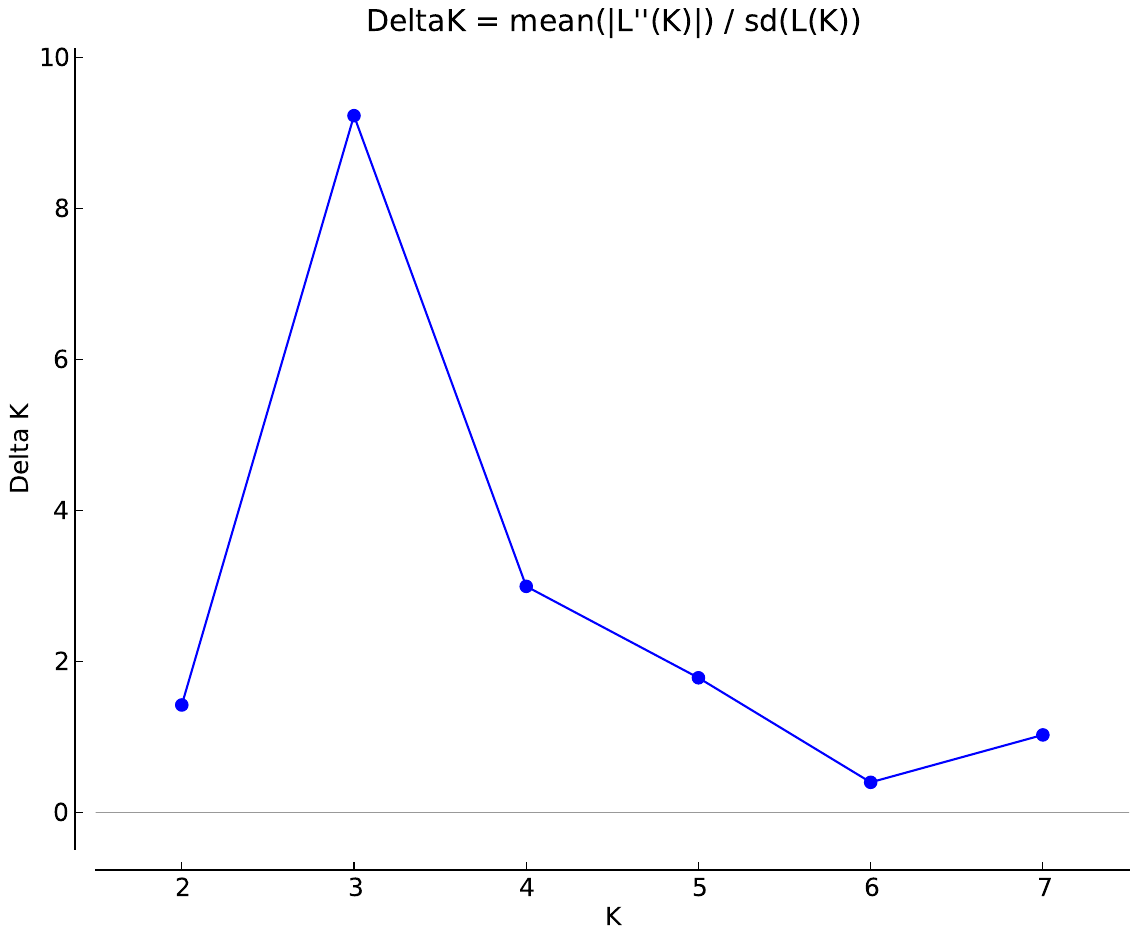

##### Figure S1. Delta-K values for the STRUCTURE runs.

Because we expect population structure to be fairly low given the recent expansion of North American starlings, we used fineRADstructure to test for more subtle patterns of structure (Malinsky et al. 2016). This program calculates shared ancestry using a coalescent model to determine haplotype linkage among sampled individuals. The resulting coancestry matrix controls for similarity among individuals to infer fine-scale patterns of population structure, and we found no evidence for subtle population structure even using this more sensitive test.

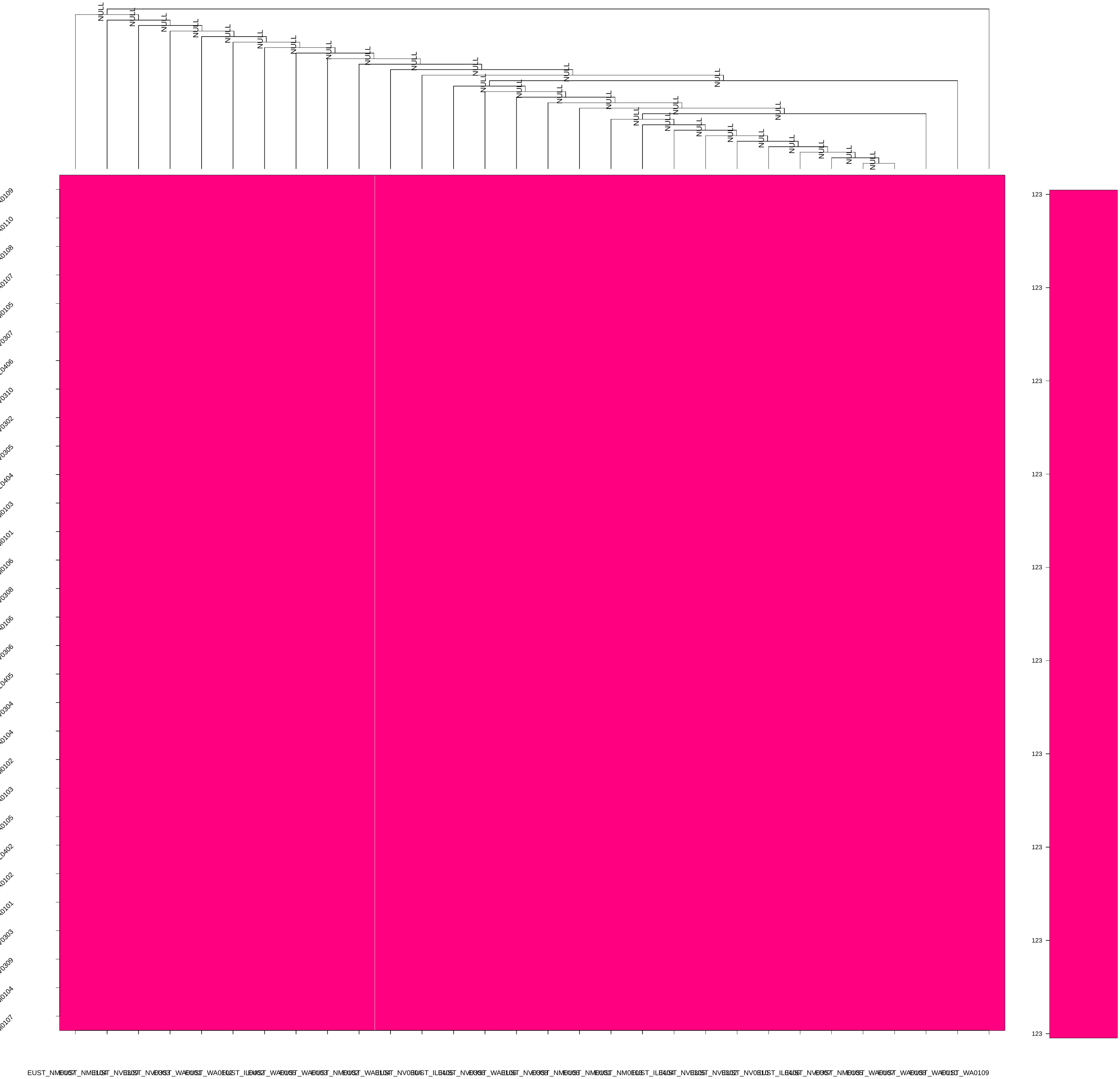

##### Figure S2. fineRADstructure results.

This clustered coancestry matrix shows that the estimated coancestry coefficient does not vary among sampled individuals. fineRADstructure identifies fine-scale patterns in population structure as a result of shared ancestry, indicating the range of coancestry coefficients using a heat map.

To identify potential geographic barriers, we also used the program EEMS (Estimated Effective Migration Surfaces, (Petkova *et al.* 2015). EEMS estimates how quickly genetic similarity decays across the landscape, allowing us to pinpoint geographic regions that depart from continuous IBD. Because the number of hypothesized demes (subpopulations) can influence model sensitivity, we ran EEMS using polygons covering the entire North American range and only the areas sampled, and also tested each map using different numbers of demes where the number of demes is limited by the number of individuals (N=50, N=100, and N=150). We adjusted the variance of the proposal distribution for both migration and diversity parameters to ensure that all parameters were accepted between 10 and 15% of the time as suggested in the EEMS documentation, with the input proposal variances as follows: mSeedsProposalS2 = 0.15, mEffctProposalS2 = 1.5, qSeedsProposalS2 = 1.5, qEffctProposalS2 = 0.1, and mrateMuProposalS2 = 0.001. We ran three chains to check convergence.

Figure S3. Estimated effective migration surfaces (EEMS) model. Warmer (red) colors indicate lower migration or diversity rates, whereas cooler (blue) colors suggest higher diversity (top) or higher migration (bottom).

The EEMS model recovers fairly uniform rates of migration among sampling locations. Although migration rates appear to be higher in some areas, these areas have not been sampled, and thus we suggest that high migration rates in this model are an artifact of sampling. Even when we decrease the number of demes, we still recover

These results, combined with previous genetic studies (Cabe 1998; 1999), support the hypothesis that the Rocky Mountains may have imposed an altitudinal barrier to starlings’ spread: spatially explicit models indicate a decreased migration rate on the eastern front of the mountains, and an increase in genetic diversity west of the mountain range (Figure 2). In other words, Western starling populations on the range edge are more diverse than populations nearer to the introduction site. Historical records complement this evidence, as the starling expansion slowed only when reaching these mountains (Jernelov 2017). We suggest that elevation may impose a barrier even across this species’ worldwide range: both in this study and in a parallel study of Australian populations, patterns of genetic variation can be attributed to a montane barrier (Cardilini 2016).

### Demographic models

The Stairway plot method estimates recent population histories from hundreds of unphased, low-coverage loci, which distinguishes the stairway plot from other demographic methods (e.g., PSMC) that can infer ancient population history more accurately (Liu & Fu 2015). The stairway plot method models changes in population size using the site frequency spectrum, where the null model assumes constant size. We used this model-flexible method to determine whether starlings experienced any genetic bottleneck after introduction: in the stairway plot, this result could occur if an alternative model was accepted during one or more steps of the stairway plot. We assumed a mutation rate of 1x10^-9^ and a generation length of 4.6 years (BirdLife International). and used the recommended 67% of sites for training. The results presented here are averaged among eight independent runs, each with 10 to 30 randomly generated breakpoints during the reconstruction.

ADD

Figure S4. Stairway plot. This model indicates a gradual decline in effective population size over time. Black line indicates approximate colonization date of starlings according to historical records (1890).

The stairway plot method finds that upon introduction approximately 130 years ago, effective population size was 10,000 individuals, and population size has gradually declined to 4,000 individuals. Importantly, the decline in N_e_ in the most recent time steps—the last 100 years—may be a spurious pattern resulting from known uncertainties in the final steps of this stairway plot method (Liu & Fu 2015).

### Rationale for GEA methods

There are several assumptions underlying the genotype-environment associations presented in this manuscript, including that: (1) RAD loci are an effective tool for exploring true genetic variation; (2) bioclimatic variation at the sampling location is a possible selective pressure that drives genetic variation in starlings; and (3) these data are best-suited to population-level inferences.

(1) We assume that RAD loci can represent actual genetic variation well enough to make inferences about population structure, demography, and even selection. Population geneticists have clearly demonstrated that the genetic diversity recovered by RAD-sequencing can be higher (CITE) or lower (CITE) than true values. In addition, RAD loci are limited to the cut sites of the enzymes used, and thus do not reflect a random sampling of genes (CITE). These realities limit the utility of RAD markers, but RAD-sequencing is nonetheless a cost-effective method for exploring genetic diversity and differentiation prior to more thorough sampling of the genome.

(2) Our study assumes that the environmental conditions experienced in the collection location represent the conditions that a bird experiences throughout its lifetime. In the main text, we present isotopic evidence that starlings in some sampling locations appear to permanently reside in or near that collection state, but here we focus on the environmental variation across time and space in our dataset. We hypothesized that conditions at the sampling location may drive selection, but environmental conditions do vary between breeding and wintering ranges. Starlings collected in the western U.S. remain in the same region during both breeding and overwintering, but elsewhere in the U.S., starlings may not experience uniform environmental conditions across their lifetime. Importantly, the environmental variation that we use in our genotype-environment associations represents an average of conditions at that location between 1970-2018 and not conditions experienced at the time of sampling. We assume that this average is a more accurate representation of potential selective pressures than a single point estimate of the conditions experienced by that bird at the time of collection.

(3) This study focuses on potential local adaptation across North America, but we acknowledge that individual dispersal and migration may complicate our ability to make inferences about selection. We cannot compare individuals’ molt origin and RAD-sequencing data side-by-side, since the original collection of starlings did not identify individuals and instead pooled samples within a population. Because we cannot trace individuals in this comparison, we cannot determine whether individual birds that carry potentially adaptive variants are more likely reside in a location year-round. Although migratory strategy can vary within the same sampling location, the molt-origin data presented in Werner et al. do enable us to determine whether individuals sampled at a given location are likely to experience consistent environmental conditions.

### Mantel tests

Why Mantel may not be appropriate for these data

As described in the main text of the Methods, we also ran partial Mantel tests control for geographic distance when testing for the relationship between environmental and genetic distances. We report all Spearman correlations here, where the P-value is the two-sided P-value after randomizing the genetic distance matrix 999 times.

##### Table S1. Partial Mantel tests.

| Variable | Definition | Mantel R | P-value |
| --- | --- | --- | --- |
| BIO1 | Annual mean temperature | -0.07 | 0.98 |
| BIO12 | Annual precipitation | 0.06 | 0.02 |
| BIO16 | Precipitation of wettest quarter | 0.28 | 0.01 |
| BIO7 | Temperature annual range | 0.15 | 0.001 |
| elevation | - | 0.15 | 0.001 |

### Comparing selection-scan methods

Differentiation methods can identify loci that have undergone strong selective sweeps, but these methods may be inappropriate in systems like this one with low overall differentiation. In this and other invasions, it is challenging to determine whether drift or selection is the major driver of differentiation. We discuss demographic evidence in the main text, but given that the effective population size expanded so rapidly after the initial founder effect, we argue that the relative importance of selection in shaping genetic variation in North American starlings may be stronger than in other invasions. However, any evidence for selection is likely to be subtle, and each of the selection tests used in this project have different assumptions that we outline in this section.

Traits under selection—especially environmentally-driven selection—are likely to be polygenic, and thus we would expect to find low levels of differentiation at many loci (CITE). For this reason, we suggest that methods such as redundancy analyses and, to a lesser degree, latent-factor mixed models may be more appropriate tests of selection in this system than differentiation-based methods.

Results from the redundancy analysis are thus reported in the main text, whereas we report the methods and results of all other tests here. Briefly, there is little overlap among these tests: no SNPs were identified by all tests, 13 SNPs were identified by both RDA and LFMM, and 3 SNPs by LFMM and BayeScEnv.

### Bayescan and BayeScEnv

F_ST_-based genome-scan approaches are best suited to identify loci that stand out against a low background level of differentiation. To test these more traditional methods against the model-based approaches (LFMM & RDA), we used both BayeScan (Foll & Gaggiotti 2008) and BayeScEnv (de Villemereuil & Gaggiotti 2015). BayeScan identified no SNPs with a Q-value lower than 0.989, indicating that there is no evidence of selection based on this test (Table S2; excel file). BayeScEnv incorporates environmental differentiation when identifying outlier loci by including a term to explicitly model environmental differentiation in the framework used in BayeScan. BayeScEnv identified several SNPs that could be under selection (Table S2), and three of these were also found the LFMM analysis (below).

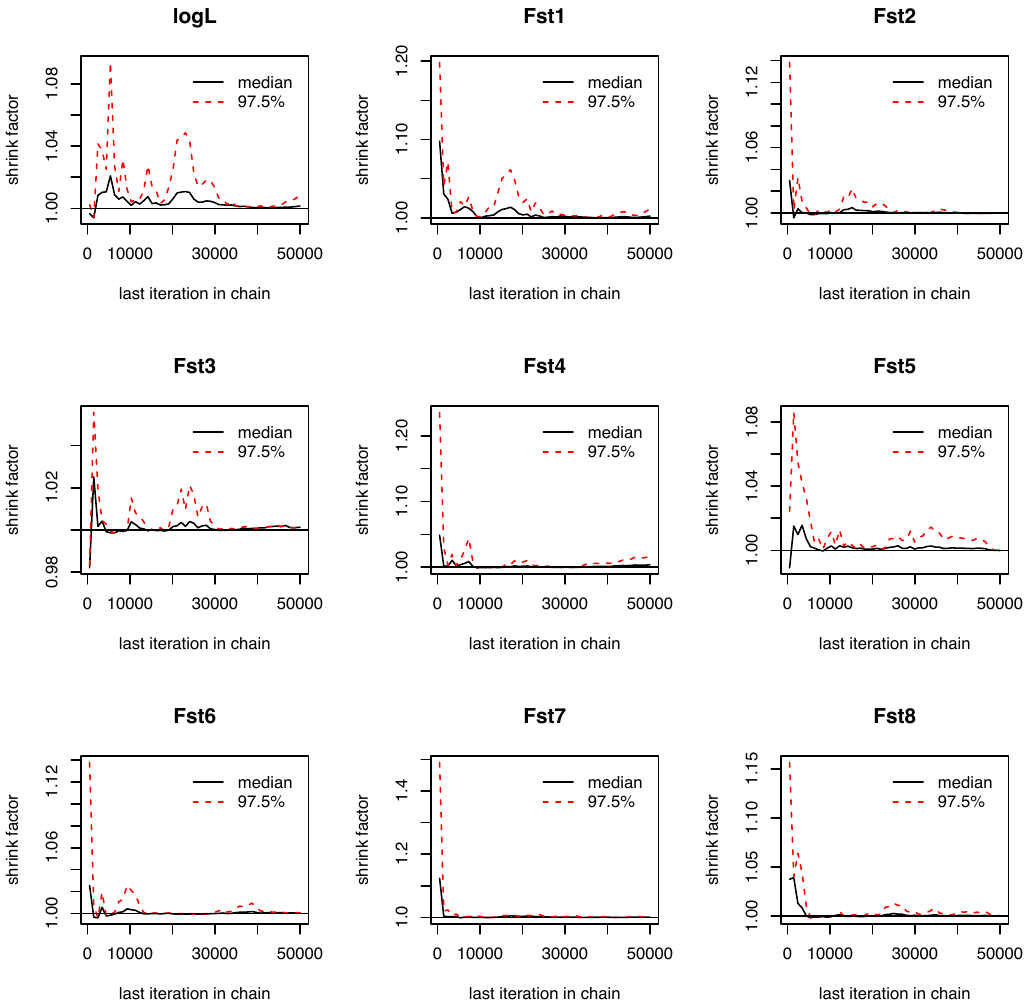

Figure S5. Gelman Plot for BayeScEnv. Here, the shrink factor demonstrates convergence of each model.

### Latent-factor mixed models (LFMM)

As a univariate test of selection, we used the *lfmm* function (Frichot *et al.* 2013) to test for associations with climatic gradients and to decrease the number of false positives. For the univariate method (LFMM), environmental variation was modeled as the first three principal components of bioclimatic variation across the range of North American starlings. We used the R package *LEA* (Frichot & François 2015) to prepare input files and run a model where genotypic variation is considered a response variable in a linear regression that controls for latent factors (e.g., population structure and/or background variation) in estimating the association between the genotypic response and the environmental predictor. For each of three models—including 1, 2, and 3 latent factors—we ran 30 MCMC chains of 10,000 cycles each, discarding a burn-in of 5,000 cycles. Z-scores were combined across all 30 runs and p-values readjusted to calibrate the null hypothesis and increase power using the Fisher-Stouffer method as suggested in the LEA and LFMM manuals. We used the Benjamini-Hochberg algorithm to control for false discoveries.

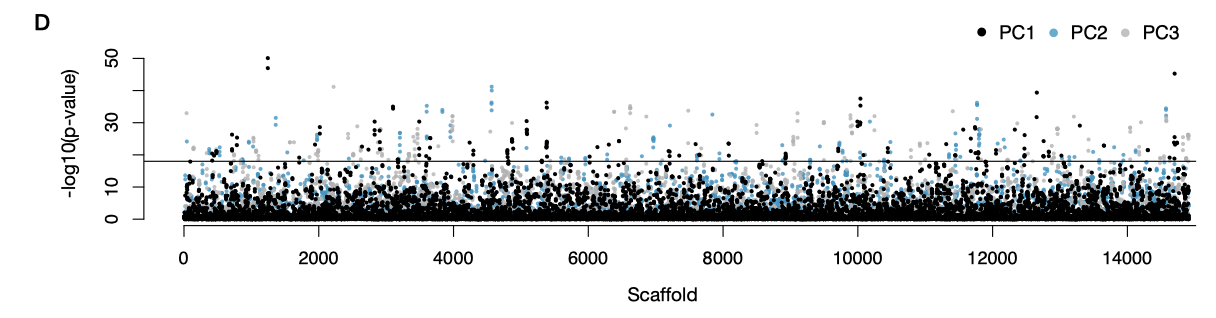

Figure S6. Latent-factor mixed model (LFMM) results. Each point reflects an association between a SNP and environmental variation (captured as a principal component of all possible bioclimatic variation across sites).

Latent-factor mixed models identified 2490 candidate variants associated with the first principal component of environmental variation, which explains 41.5% of the variation and loads with temperature-related variables. An additional 1315 variants were associated with precipitation-related PC2, and/or with PC3, a composite of temperature and precipitation variables. Since we identified many candidates using a q-value cut-off of 0.01, only loci that were identified in all three runs (K=1-3 latent factors) and were more than five standard deviations from the mean log10p value were considered candidates under selection (false discovery rate < 0.05, true positive >25). This filtering left 1218 remaining SNPs—or 8% of all SNPs—distributed across all three principal components of environmental variation (Table S2).

### Functions of genes and gene ontology information

Although no gene ontology categories were significantly overrepresented, signaling and response to stimuli were particularly well-represented among GO terms, showing up to 48-fold enrichment (FDR-corrected P=0.12-0.98, Table S2). It is important to note that this analysis does not correct for gene size nor does it expect that any ‘candidates’ reported here are likely to drive adaptation in the North American starling. However, we find it useful to examine possible functions that would need to be verified by whole-genome data. Among signaling-related GO terms, neuron development, synaptic transmission and organization were particularly common (Table S2). Other common GO terms relate to kidney function, viral processing, metabolism, and regulation of growth factors. Among the top twenty variants under strong selection (*r*^2^ > 0.2 or log_10_*p* > 10), we find four genes related to growth factors (EOGT, GAB3, HBEGF, STAT3), six involved in immune responses that do not directly involve growth factors (DNAJB14, FKBP4, ASB2), and three essential to muscle function (LIMCH1, HBEGF, CALD1). Putatively selected genes may play a role in physiological processes that support starlings’ invasion success in North America.

Effective solute transport and kidney development are critical in dry habitats, which may explain why all but one of the genes related to kidney function correlate with precipitation (BIO16). Claudin 16 (CLDN16; *R*^2^ = 0.23) is one such protein that regulates ion concentrations in the kidney, while others maintain homeostasis and vasoconstriction (AVPR1B; *R*^2^ = 0.18) or transport iron (STEAP3; *R*^2^ = 0.21). Many invaders shift their diet upon colonization of a new habitat, and many candidates play a role in metabolism and/or digestion: for example, aridity may result in selection on proteins that process lipids (MTMR3; *R*^2^ = 0.18) and fatty acids (PEX5; *R*^2^ = 0.17), since organisms living in dry environments often depend on fat storage for proper hydration. Proteins that modify growth factors—key orchestrators of cellular growth and development—correlate with aridity but also temperature, and complexes that rely on ubiquitin ligases to degrade proteins are similarly strong candidates. Many of the putatively selected genes are critical to starlings’ survival, and investigating a wider range of environmental conditions and sampling whole genomes may support this preliminary evidence of incipient local adaptation.
